# aFARP-ChIP-seq, a convenient and reliable method for genome profiling in as few as 100 cells with a capability for multiplexing ChIP-seq

**DOI:** 10.1101/474676

**Authors:** Wenbin Liu, Sibiao Yue, Xiaobin Zheng, Jia Cao, Yixian Zheng

**Author notes:** Correspondence (XZ), (JC), and (YZ).

## Abstract

Much effort has been devoted to understand how chromatin modification regulates development and disease. Despite recent progress, however, it remains difficult to achieve high sensitivity and reliability of chromatin-immunoprecipitation-coupled deep sequencing (ChIP-seq) to map the epigenome and global transcription factor binding sites in cell populations of low cell abundance. We present a new Atlantis dsDNase-based technology, aFARP-ChIP-seq, that provides accurate profiling of genome-wide histone modifications in as few as 100 cells. By mapping histone lysine trimethylation (H3K4me3) and H3K27Ac in group I innate lymphoid cells from different tissues, aFARP-ChIP-seq uncovers potentially distinct active promoter and enhancer landscapes of several tissue-specific NK and ILC1. aFARP-ChIP-seq is also highly effective in mapping transcription factor binding sites in small number of cells. Since aFARP-ChIP-seq offers reproducible DNA fragmentation, it should allow multiplexing ChIP-seq of both histone modifications and transcription factor binding sites for low cell samples.

## Introduction

Chromatin immunoprecipitation coupled with deep sequencing (ChIP-seq) is a powerful technique for genome-wide mapping of the binding of chromatin regulators and epigenetic modifications, which has contributed greatly to both basic and translational research (Park, 2009; Furey, 2012). For example, the accurate mapping of epigenome changes in cell populations at distinct developmental stages facilitate our understanding of epigenetic mechanisms by which different cell lineages establish their unique transcriptional programs. Unfortunately, two major limitations restrict the utility of conventional ChIP-seq method in studying rare cell types isolated directly from tissues. The first is fragmentation. Although sonication is the most commonly used approach for chromatin fragmentation in ChIP-seq, it can result in epitope damage and thus reducing the immunoprecipitation efficiency especially when the initial material is limited (Stathopulos *et al.*, 2004). The inconsistencies in sonication-based chromatin fragmentation results in a low-throughput processing of samples because each sample needs to be tested for specific setting of sonication power and time. This impedes its adaptation for reliably processing of multiple samples. Although micrococcal nuclease (MNase) has been used as an alternative to sonication for chromatin fragmentation, MNase often causes chromatin over-digestion (Brind’Amour *et al.*, 2015). Another difficulty is chromatin loss during multiple steps of ChIP-seq operation, which makes it difficult to obtain high-quality mapping in a small number of cells (Park, 2009).

Several strategies have been employed to reduce the number of cells needed to produce high quality ChIP-seq. One method is based on increasing DNA amplification cycles during the sequencing library building, which allowed high quality ChIP-seq in thousands of cells (Adli *et al.*, 2010; Adli and Bernstein, 2011; Shankaranarayanan *et al.*, 2011; Ng *et al.*, 2013). The major deficiency of the amplification-based method is that the low-abundance of some ChIPed chromatin may be underrepresented or lost, which makes the method not applicable to ultralow cell numbers. Another method is barcoding of the fragmented chromatin in individual samples followed by sample pooling (Lara-Astiaso *et al.*, 2014; Rotem *et al.*, 2015; van Galen *et al.*, 2016). By pooling multiple barcoded samples, the increased total ChIPed chromatin helps to reduce chromatin loss in subsequent steps. The low efficiency of ligating barcode adaptors to chromatin fragments (<10%), however, significantly limits the application of the method for low cell numbers (Gury-BenAri *et al.*, 2016b), because the majority of chromatin failed to be barcoded. Indeed, between 10,000 to 20,000 sorted hematopoietic cells were needed for the barcode-based iChIP-seq studies (Lara-Astiaso *et al.*, 2014).

By combining microfluidics and DNA barcoding, Rotam et al reported the mapping of chromatin states at single-cell resolution and identified a spectrum of heterogeneity defined by differences in chromatin signatures of pluripotency and differentiation priming among mouse embryonic stem cells (mESCs) (Rotem *et al.*, 2015). While this single-cell method is useful for identifying subpopulations of cells with the aid of single-cell RNA-seq data, the low number of valid sequencing reads (500 −10,000) per cell makes the method insufficient for de novo regulatory site identification (Rotem *et al.*, 2015). Another recently published method, ChIPmentation, utilizes Tn5 transposon mediated adapter addition to immunoprecipitated chromatin still bound to beads. This greatly simplified the ChIP procedure, thereby reducing the time and cell input requirement (Schmidl *et al.*, 2015). However, since chromatin loss still occurs during the sonication step, ChIPmentation requires ~10,000 cells for accurate mapping of histone modifications of H3K4me3 and H3K27me3.

Innate lymphoid cells (ILCs) are a family of recently defined lymphoid cells that belongs to the innate counterpart of lymphocytes (Rankin *et al.*, 2013; Diefenbach *et al.*, 2014; Eberl *et al.*, 2015). ILCs are present throughout the body and they are often found at the barrier surfaces of tissues. These cells play important roles in early defense against pathogens and they promote tissue repair and maintenance. Their inappropriate activation could contribute to inflammation and autoimmune diseases (Rankin *et al.*, 2013; McKenzie *et al.*, 2014). Developing from the common ILC progenitor, three classes of ILCs have been defined, including group 1, group 2, and group 3 ILCs. These ILCs are characterized based on their analogous cytokine-production profiles of the adaptive T cell subsets (Diefenbach *et al.*, 2014). The group 1 ILCs consist of conventional Natural Killer cell (NK) and ILC1. Controversial classification exists in this subgrouping because they have functional similarities and shared expression of many cell surface markers (Jiao *et al.*, 2016). Recent global transcriptome profiling of group 1 ILCs isolated from different peripheral tissues has begun to facilitate the subdivision of the NK and ILC1 subsets (Robinette *et al.*, 2015). For example, the small intestine intraepithelial (siIEL) ILC1 is found as a unique subset with distinct developmental and functional properties (Fuchs *et al.*, 2013). The unique transcription and cytokine profiles suggest that siIEL ILC1 may belong to a distinct ILC1 subset different from some of the other ILC1 or NK subsets (Robinette *et al.*, 2015). However, the lack of high quality epigenome and chromatin protein binding profiles due to the difficult to perform ChIP-seq in the low abundance siIEL ILC1 has limited additional in-depth study of the lineage identity and origin of these cells (Sciumè *et al.*, 2017).

We have recently report two techniques, recovery via protection (RP)-ChIP-seq and favored amplification RP-ChIP-seq (FARP-ChIP-seq) for low-cell-number epigenome profiling (Zheng *et al.*, 2015). These two ChIP-seq methods are based on the idea that if a rare cell population is not lost during the initial fixation and wash steps, and if chromatin loss is minimized during ChIP and library building, it should be possible to recover low-abundance chromatin without increasing DNA amplification cycles. Indeed, by employing these two methods, we were able to obtain reproducible mapping in as few as 500 cells. Here, we report a new dsDNase-based FARP-ChIP-seq method for high-fidelity genome-wide profiling using as few as 100 cells with broad applications such as profiling of epigenetic differences of group 1 ILCs from different tissue origins and transcription factor binding in splenic B cells. The reliability and consistency of dsDNase-based chromatin fragmentation among different samples in our method also allows multiplexing of ChIP-seq operations.

## Results

### Atlantis dsDNase-based FARP-ChIP-seq (aFARP-ChIP-seq) offers better mapping of histone marks than FARP-ChIP-seq

We recently reported RP-ChIP-seq and FARP-ChIP-seq that allow high quality epigenome mapping in only 500 cells. However, the increase in sequencing depth required due to the presence of carrier DNA leads to an increase in mapping costs (Zheng *et al.*, 2015). To reduce both the sequencing costs and the number of cells needed for successful FARP-ChIP-seq, we wish to further improve the recovery of chromatin of interest. We reasoned that sonication used for chromatin fragmentation could destroy the epitopes for subsequent immunoprecipitation, thereby resulting in a significant loss of chromatin of interest especially when the input cell number is low. To overcome this limitation, we tested enzymatic digestion-based chromatin fragmentation. We found that the commonly used micrococcal nuclease (MNase)-based fragmentation is compatible for FARP-ChIP-seq and it resulted in ~70% increase (from 16% to 27%) of H3K4me3 reads mapping to mouse genome compared to FARP-ChIP-seq at the same read depth for 500 mESCs (Fig 1A and B). However, it is challenging to control MNase activity and its propensity to over digest chromatin due to its exonuclease activity impedes reliable parallel processing of multiple samples, especially when dealing with different samples (Brind’Amour *et al.*, 2015; van Galen *et al.*, 2016). Therefore, we sought to identify an alternative DNase for chromatin fragmentation.

**Figure 1.**
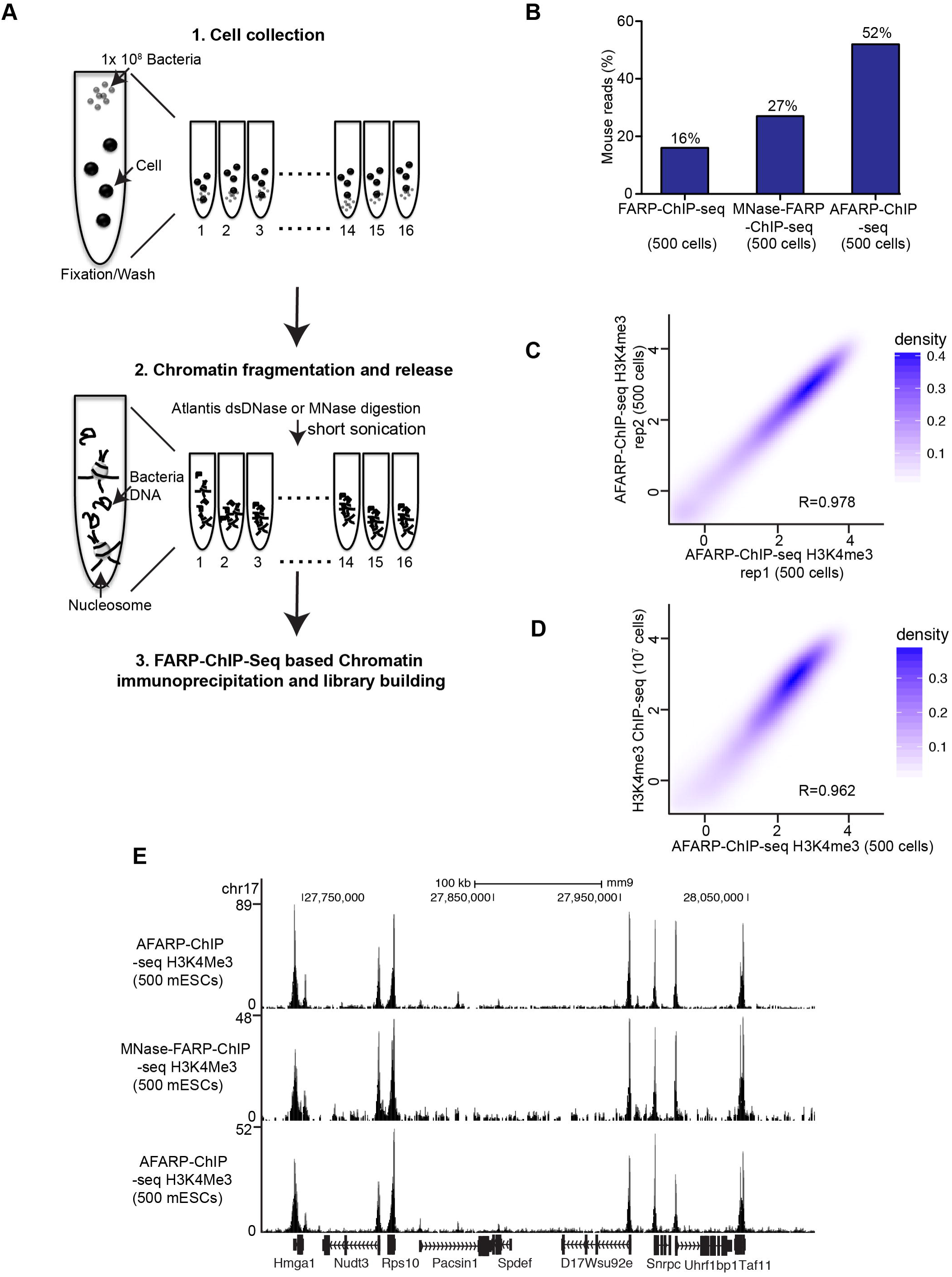
Development of aFARP-ChIP-seq. (A) Schematic overview of aFARP-ChIP-seq or MNase-FARP-ChIP-seq. For aFARP-ChIP-seq, 1-16 samples can be simultaneously processed with consistent quality due to the consistent chromatin fragmentation in different samples. (B) aFARP-ChIP-seq resulted in increased ratios of mapped mouse reads to the total reads (mouse reads+carrier biotin-DNA+bacteria DNA reads). (C and D) Contour plots (as log2 of the average read density within 2 kb up and downstream of TSS. Spearman correlation coefficient, R) between two biological replicates of aFARP-ChIP-seq in 500 mESCs (C) or between 500 mESC AFARP-ChIP-seq and the 10^7^ mESCs standard ChIP-seq of H3K4me3. (E) Enrichment at the indicated genes on chromosome 17 was mapped by the indicated methods.

See also Figure S1.

Among the DNases, the Atlantis dsDNase (Zymoresearch) is a double-stranded DNA-specific endonuclease that cleaves phosphodiester bonds in DNA to yield fragments with 5’-phosphate and 3’-hydroxyl termini, which are ideal for chromatin fragmentation. Our tests demonstrated that Atlantis dsDNase digestion of unfixed or paraformaldehyde fixed nuclei from different cell types and cell numbers at 0.5 Unit (U) for 20-30 min at 37 °C yielded consistent chromatin fragmentation (Figure S1 and data not shown). This reliable fragmentation and suitable DNA length distribution are optimal for ChIP-seq (Park, 2009) and it is amenable for simultaneously fragment chromatin in different samples (Figure 1A). Thus, the Atlantis dsDNase may replace the conventional MNase for chromatin fragmentation in a variety of genome studies.

We next attempted at incorporating Atlantis dsDNase for chromatin fragmentation in our FARP-ChIP-seq and we referred to this method as Atlantis-FARP-ChIP-seq (aFARP-ChIP-seq). Since the total chromatin marked by H3K4me3 is much lower compared to other histone modifications, it is most challenging in obtaining sufficient chromatin for high quality H3K4me3 profiling using low cell numbers. Thus, we initially applied aFARP-ChIP-seq to map H3K4me3 in 500 mESCs (Figure 1A; also see the method section). We found that aFARP-ChIP-seq resulted in a ~3-fold increase of DNA reads of interest compared to FARP-ChIP-seq at the same read depth for 500 mESCs (Figure 1B). The two biological replicates were highly consistent (Figure 1C), demonstrating the reproducibility of this method. To further evaluate aFARP-ChIP-seq, we performed genome-wide comparisons of H3K4me3 signal intensity against the datasets generated from the standard ChIP-seq of 10^7^ mESCs (Jia *et al.*, 2012). This revealed that aFARP-ChIP-seq using 500 mESCs was highly correlated with the standard ChIP-seq of 10^7^ mESCs (Figure 1D). Analyses of specific chromatin regions revealed that aFARP-ChIP-seq could uncover H3K4me3 peaks reliably. More importantly, aFARP-ChIP-seq generated higher H3K4me3 signal intensity compared to FARP-ChIP-seq (Figure 1E). Thus, aFARP-ChIP-seq yields accurate and consistent epigenome profiles and it performs better compared to FARP-ChIP-seq relying on the sonication-based or MNase-based chromatin fragmentation in 500 cells (Figure 1B).

### aFARP-ChIP-seq epigenome mapping in as few as 100 cells

The improved mapping efficiency of aFARP-ChIP-seq suggests that it could be used in less than 500 cells. To test this, we mapped H3K4me3 using 100 mESCs. We digested the fixed mESCs for 20 min. We found that the H3K4me3 mapping in 100 mESCs showed good consistency between the two biological replicates (Figure 2A). Importantly, despite increased noise, the chromatin profiles generated from 100 mESCs were still informative and were similar to the 500-cell aFARP-ChIP-seq (Figure 2B). By using the MACS2 program with identical parameters, the peaks called for our previous standard ChIP-seq of 10^7^ mESCs datasets (Jia *et al.*, 2012) and aFARP-ChIP-seq of 100 mESCs also have a good degree of overlap (Figure 2C).

**Figure 2.**
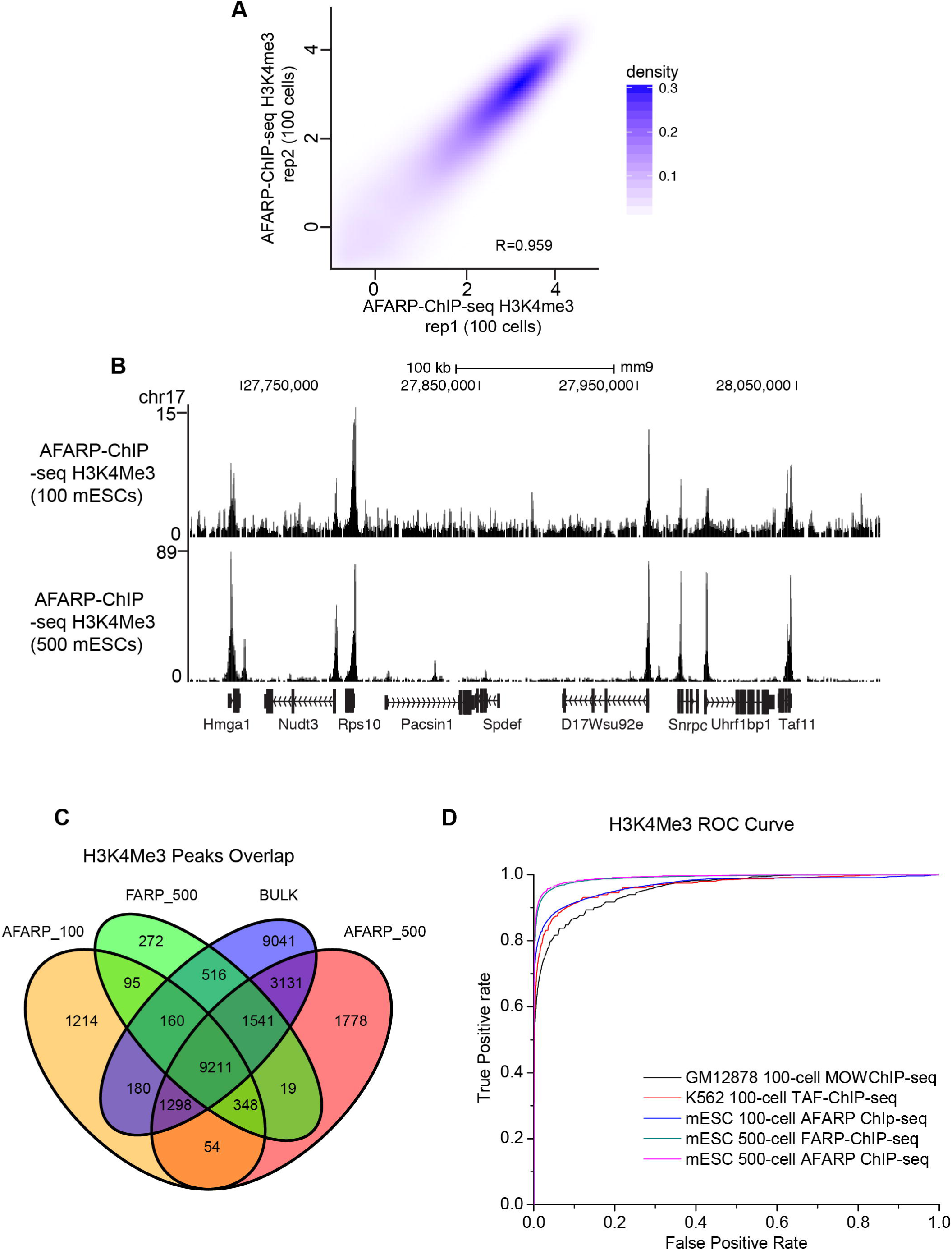
aFARP-ChIP-seq generates reliable epigenome mapping in as few as 100 cells. (A) Contour plots between two biological replicates of aFARP-ChIP-seq of H3K4me3 in 100 mESCs. (B) Enrichment at the indicated genes on chromosome 17 was mapped by the aFARP-ChIP-seq with 100 or 500 mESCs. (C) The overlap between the H3K4me3 peaks identified from conventional ChIP-seq of 10^7^ mESCs and FARP-ChIP-seq of 500 mESCs and aFARP-ChIP-seq of 100 and 500 mESCs. MACS2 with identical parameters was used to identify the peaks. (D) ROC curve for H3K4me3 comparisons among different cell number inputs and ChIP-seq methods. True positive rate, the number of true-positive 2-kb regions divided by the number of ‘‘true’’ regions (25,000); false-positive rate, the number of false positive regions divided by the number of ‘‘false’’ regions.

Next, we used the receiver-operating characteristic (ROC) curve to compare the H3K4me3 maps obtained by FARP-ChIP-seq or aFARP-ChIP-seq. The Area Under the ROC curve (AUC) is a standard metric for quantifying balanced sensitivity and specificity. By using different cutoffs to calculate the true-positive and false-positive rates, we plotted ROC curves for each method, which showed that aFARP-ChIP-seq using 100 or 500 cells provides reliable performances (Figure 2D). These analyses demonstrate that aFARP-ChIP-seq enables analysis of as few as 100 cells.

### The H3K4me3 profiling of group 1 innate lymphocyte lineages reflects their distinct functionalities

To test the applicability of aFARP-ChIP-seq, we applied it on challenging *in vivo* biological samples by profiling histone modifications in innate lymphoid cell (ILC) types, which typically make up 1-5 % of total lymphocytes in peripheral non-gut tissues (Diefenbach *et al.*, 2014). Recent studies indicate that tissue-specific signals have significant impacts on gene expression and activity of ILCs. The influence of local tissue microenvironments on chromatin and gene regulatory landscapes, however, has remained not well understood (Gury-BenAri *et al.*, 2016a; Shih *et al.*, 2016; Sciumè *et al.*, 2017), in part, due to the difficulty in mapping chromatin and gene regulatory landscapes in ILCs isolated from individual animals. We focused our study on the IFN-γ-producing group 1 ILCs, including conventional NK and ILC1, isolated from individual mice (Cortez and Colonna, 2016)

Since there is a lack of unique and consistent markers in NK cells and ILC1 cells in various organs, we used different sorting strategies according to previously published protocols for each tissue, including spleen, mesenteric lymph nodes (mLN), liver and small intestine intraepithelia (Figure 3A and Figure S2A) (Robinette *et al.*, 2015). We then applied aFARP-ChIP-seq to profile H3K4me3 using NK or ILC1 sorted from each tissue from one mice without further *in vitro* culturing. For each aFARP-ChIP-seq analyses, we used 1000-2000 cells that were estimated based on the cell numbers sorted by Fluorescence Activated Cell Sorting (FACS). We found consistent maps for H3K4me3 (Figure S2B-H) between biological replicates in the ILC1 and NK cells in different tissues, which allowed us to examine the chromatin landscapes in these two group 1 ILC sub-lineages.

**Figure 3.**
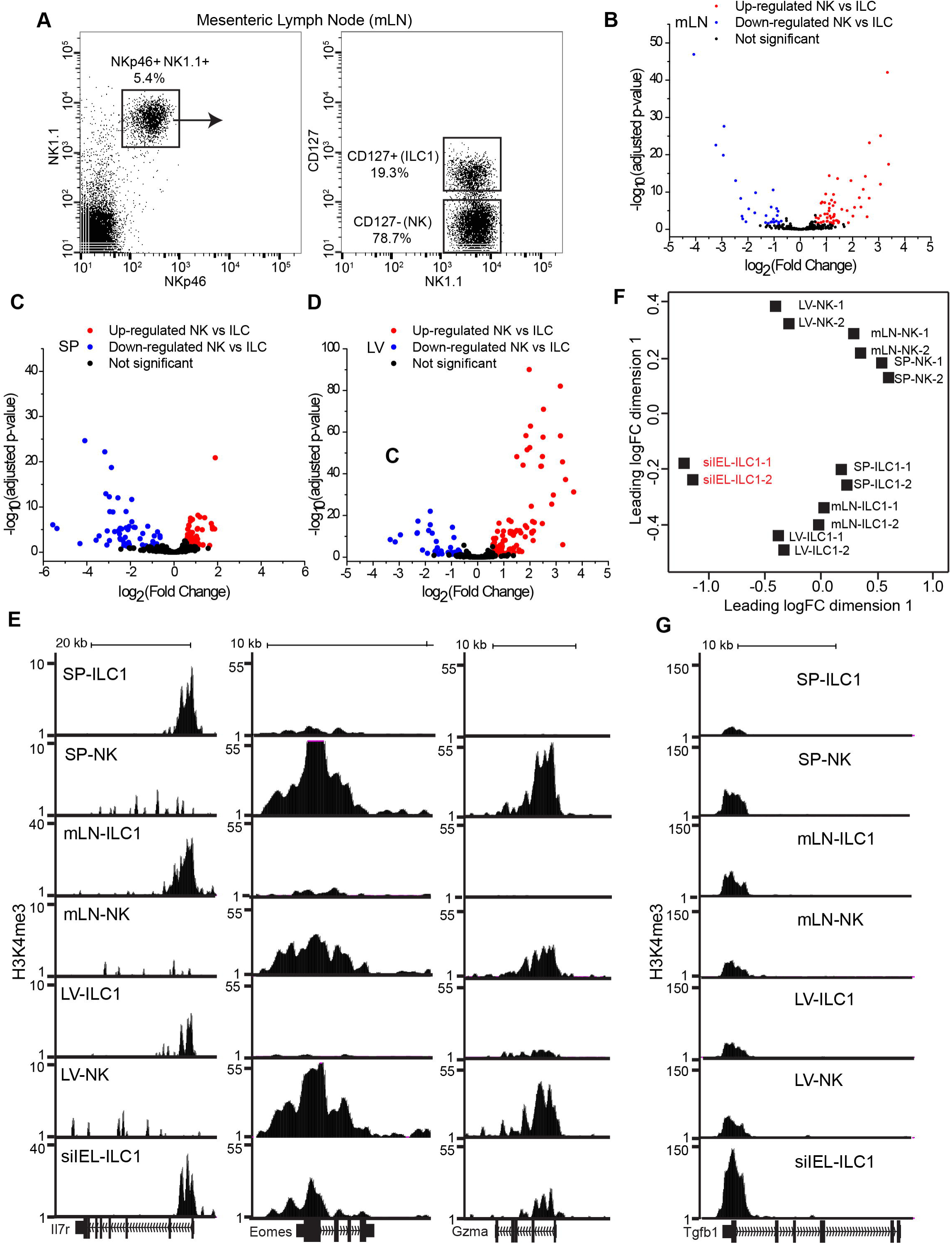
aFARP-ChIP-seq of H3K4me3 in group 1 ILCs from indicated tissues. (A) FACS plots with the markers and gating used for sorting NK and ILC1 cells from mesenteric lymph nodes (mLN). (B-D) Comparison of H3K4me3 in NK and ILC1 cells isolated from mesenteric lymph node (mLN) (B), spleen (SP) (C), and liver (LV) (D). (E) Genome browser view of the representative lineage- and function-specific H3K4me3 peaks in NK and ILC1 cells isolated from the indicated tissues. (F) MDS plot of aFARP-ChIP-seq of H3K4me3 maps in NK and ILC1 cells isolated from the indicated tissues. Distances between biological replicates (−1 and −2) within and between group 1 ILC subtypes show the similarities and differences among the datasets. (G) Browser view of TGF-β H3K4me3 peaks in NK and ILC1 from the indicated tissues.

See also Figure S2 and Table S1

We identified 789 up- and 271 down-regulated H3K4me3 peaks in spleen, 604 up- and 557 down-regulated peaks in mLN, and 966 up- and 490 down-regulated H3K4me3 peaks in liver in ILC1 compared to NK cells (fold change > 1.5, FDR < 0.05.) (Figure 3B-D). We found those exhibiting differentially H3K4me3 peaks correlated with the lineage-specific developmental program and functionality of each subset such as IL7r (ILC1), Eomes (NK), and Gzma (NK, cytotoxic machinery) (Figure 3E).

Specifically, the H3K4me3 levels on Eomes is greater than twofold in NK cells than in ILC1 cells in all tissues analyzed (Figure 3E and Table S1), which is consistent with previous finding that Emoes is a marker for NK cells (Gordon *et al.*, 2012; Daussy *et al.*, 2014; Zhang *et al.*, 2018). Interestingly, we found that siIEL ILC1 isolated from small intestines exhibited H3K4me3 peaks on both IL7r and Eomes (Figure 3E). This suggests functional plasticity of this unique ILC1 probably due to their constant exposure to varied environmental signals from microbiome and nutrients in the gut. The Multi-Dimensional Scaling (MDS) plot also revealed that siIEL ILC1 to be distinct from both ILC1 and NK derived from different peripheral tissues (Figure 3F). The elevated H3K4me3 peak in TGF-β locus in siIEL ILC1 compared to NK and other ILC1 cells (Figure 3G) is consistent with the unique roles of TGF-β in the development and function of siIEL ILC1 (Robinette *et al.*, 2015; Cortez *et al.*, 2016)

### H3K27Ac mapping reveals differential enhancer landscapes in the siIEL ILC1 compared to the other ILC1

It is well known that gene expression programs in ILC subsets in different tissues can reflect distinct patterns of enhancer activity that in turn reflects the differential transcription factor binding profiles (Hallikas *et al.*, 2006; Spitz and Furlong, 2012; Heinz *et al.*, 2015). To probe the regulatory circuitry that specifies siIEL ILC1 and its functions, we next mapped the active enhancer mark H3K27Ac (Creyghton *et al.*, 2010) in the same set of NK and ILC1 isolated from individual mice as described above (see Figure 3 and S2) using estimated 1000-2000 cells (based on FACS sorting) for each ChIP-seq experiment. The biological replicates of our maps were highly consistent with one another (Figure 4A and Figure S3A-F). We then focused on analyzing the active enhancers in ILC1 from spleen, lymph nodes, liver, and small intestine. We identified total 21417 enhancers that are active in at least one ILC1 cell type from at least one of the four tissues profiled (Figure 4B and Table S2). Consistent with previous finding of an early developmental acquisition of common chromatin organization in ILCs (Shih *et al.*, 2016), the ILC1 cells from siIEL shared 9892 enhancers with the other ILC1 from spleen, lymph node, and liver (Figure 4B). We also identified 2100 enhancers that are unique to siIEL ILC1 (Figure 4B).

**Figure 4.**
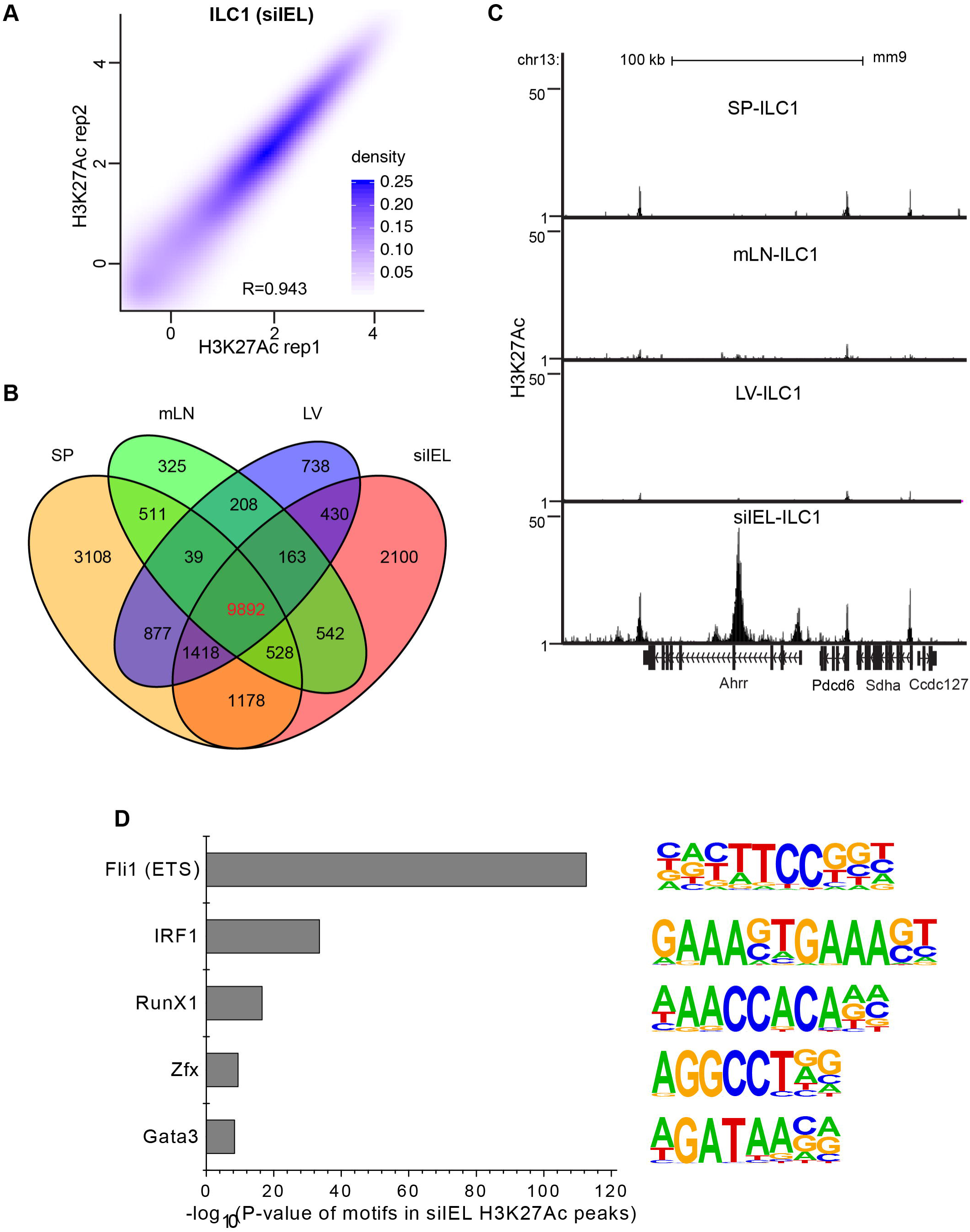
Identification of the unique enhancer landscape in siIEL ILC1 among the four ILC1s mapped. (A) Contour plots between two biological replicates of aFARP-ChIP-seq of H3K27Ac in siIEL ILC1. (B) The Venn diagram showing the numbers of active enhancers that are either unique or shared by the indicated number of the indicated tissues. (C) Genome browser view of H3K27Ac map at and around Ahrr locus in ILC1s from the indicated tissues. (D) P values for the enrichment of transcription factor binding motifs identified by analyzing enhancers that are active in siIEL ILC1. The sequences of the motifs are listed to the right.

See also Figure S3 and Table S2

By analyzing the top five genes located most proximally to the up-regulated enhancers in the siEIL ILC1 (compared to the other three ILC1), we found Ahrr, Cnih3 (Figure 4C and Figure S3G), Ccny, Sec24d, and Rin2. Interestingly, a recent study reported that the expression of AhRR in colonic intraepithelial lymphocytes prevents excessive IL-1β production and Th17/Tc17 differentiation, implicating the physiologic importance of AhRR in balancing intestine inflammation (Brandstätter *et al.*, 2016). Our finding that Ahrr gene is marked strongly by active enhancers in siIEL ILC1 suggests that siEIL ILC1 could also use Ahrr in modulating inflammation in the small intestines. Our analyses also indicate that the other genes, such as Cnih3 and Ccny, could also play important roles in siEIL ILC1. Although additional studies are required to validate this possibility, the high-quality enhancer profiling achieved using aFARP-ChIP-seq in small number of cells directly isolated from tissues should facilitate the identification of candidate genes that function in lineage specification or functional plasticity of cells *in vivo* such as the ILC1 subsets in different tissue microenvironments in individual mice.

Since transcription factor binding is a key determinant of enhancer activity, we next attempted to identify potential transcription factors that could regulate the enhancer landscape in siEIL ILC1 by searching for the enrichment of transcription factor binding motifs in H3K27Ac peaks identified in these cells. This allowed the identified a full set of transcription factor signatures of siEIL ILC1. Among these, we found significant enrichment of sequence motifs known to be bound by Fli1 (or ETS), IRF1, RunX1, Zfx and Gata3 (Figure 4D), which have been shown to play important roles for the development and function of ILC1 in general (Rankin *et al.*, 2013; Diefenbach *et al.*, 2014; Tanriver and Diefenbach, 2014). These results suggest that the shared transcriptional regulatory elements underlying either development or functionality of ILC1 subpopulations across the different tissue origins. Together, our analyses show that the high quality H3K4me3 and H3K27Ac datasets we generated for NK and ILC1 in different tissues can serve as valuable resources.

### aFARP-ChIP-seq is applicable for high quality mapping of transcription factor binding sites in small number of cells

Genome-wide profiles of transcription factor binding sites have been largely restricted to tissue culture cell lines due to the requirement of a large number of cells (>10^6^ cells) as the starting material for the traditional ChIP-seq approach (Valouev *et al.*, 2008; Ouyang *et al.*, 2009). To test if aFARP-ChIP-seq can facilitate the mapping of transcription factor binding sites in relatively small number of cells sorted from tissues, we mapped the binding sites of the ETS-family transcription factor, PU.1. PU.1 is widely expressed in hematopoietic lineages and plays key regulatory roles in early hematopoiesis and B-cell development (Klemsz *et al.*, 1990). As proof of principle, we performed genome-wide mapping of PU.1 in splenic B cells using 1×10^5^ or 1×10^4^ cells in each aFARP-ChIP-seq. Several lines of evidence suggest that our profiling identified bona fide PU.1 binding sites in the isolated B cells. First of all, our data showed proper read distributions around genomic loci of genes that have previously been reported to be bound by PU.1 in B cells, such as Blnk (encoding B cell linker protein), Btk (encoding Bruton tyrosine kinase), and Fcgr2b (encoding FcγRIIb), and our parallel mapping using isolated T cells showed that these genes do not have PU.1 binding as expected (Figure 5A and Figure S4) (Schweitzer *et al.*, 2006; Xu *et al.*, 2012; Solomon *et al.*, 2015). Additionally, we observed a similar PU.1 binding pattern around TNF locus as those obtained using iChIP-seq in 1x 10^4^ dendritic cells derived by *in vitro* differentiation of mouse bone marrow cells (Figure 5B) (Lara-Astiaso *et al.*, 2014).

**Figure 5.**
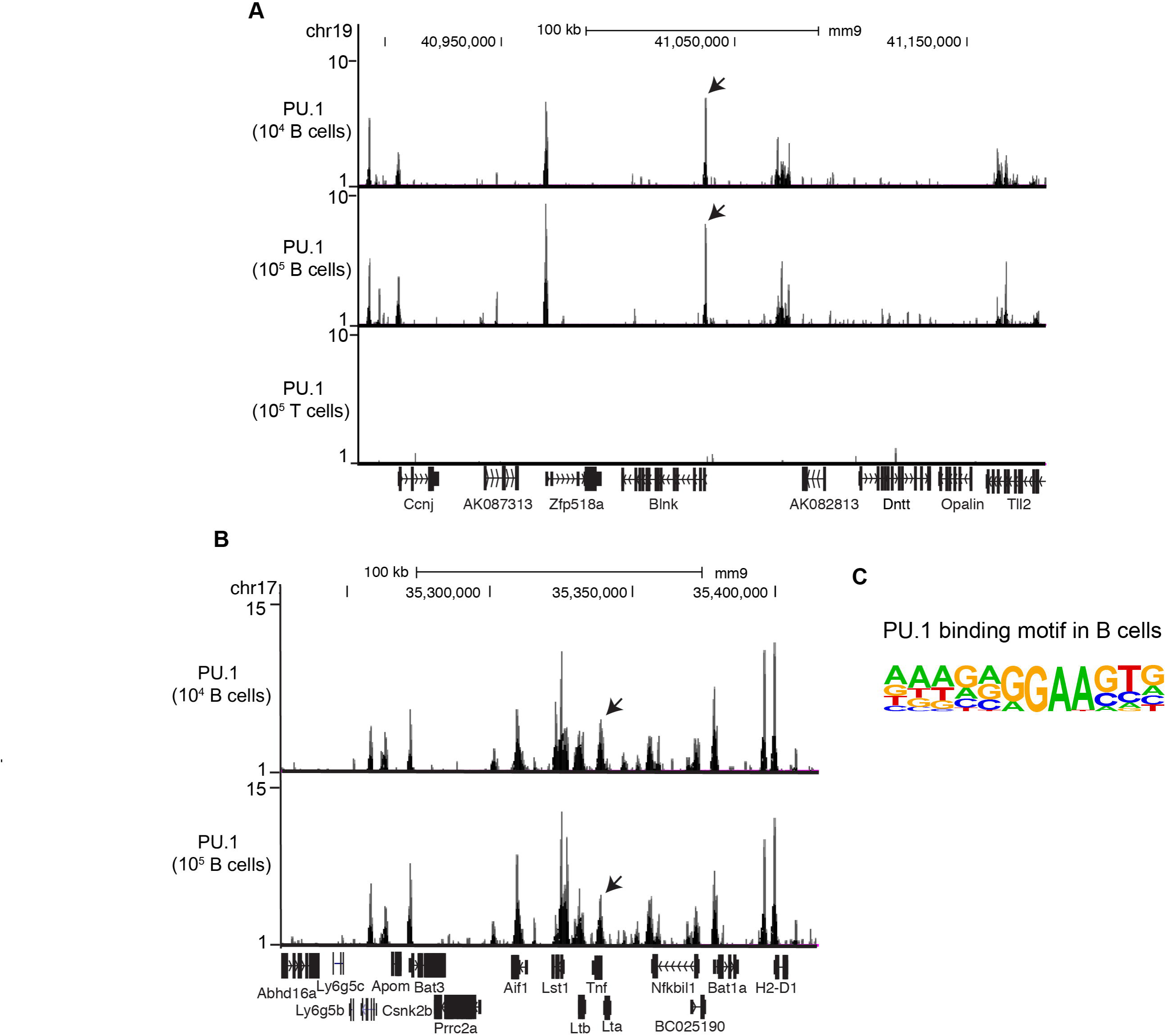
aFARP-ChIP-seq mapping of genome-wide PU.1 binding in isolated splenic B cells. (A) Normalized profiles of PU.1 in a 200 kb region surrounding the Blnk locus obtained with aFARP-ChIP-seq in splenic B and T cells. The number of cells in the input samples is indicated on the y axis. Arrows point to the peaks at the Blnk locus. (B) Normalized profiles of PU.1 in a 150 kb region surrounding the TNF locus obtained with aFARP-ChIP-seq in the indicated number of isolated splenic B cells. Arrows mark the peaks at the TNF locus. (C) The PU.1 binding motif shown was identified as the top hits based on the motif analysis of all PU.1 binding regions.

See also Figure S4

To further validate the results of our aFARP-ChIP-seq in the isolated splenic B cells, we screened for the sequences of the enriched read regions for potential transcription factor binding motifs. We found that the top motif for PU.1-bound regions contain a canonical ETS motif, 5′-GGAA-3′ (Figure 5C) (Solomon *et al.*, 2015). Therefore, the reproducibility and sensitivity of aFARP-ChIP-seq should allow genome wide characterization of chromatin binding proteins including transcription factors in small number of cells directly sorted from tissues.

### Discussion

By searching for DNases that allow reliable chromatin fragmentation under wider range of conditions than the commonly used MNase, we found that the Atlantis dsDNase can fragment chromatin reproducibly in different cell types and in different number of cells using a relatively wide range of incubation time without over digesting chromatin. Thus, the Atlantis dsDNase is a superior choice over MNase in applications involving chromatin fragmentation.

By applying FARP-ChIP-seq on the Atlantis dsDNase generated chromatin (aFARP-ChIP-seq), we are able to produce high-quality ChIP-seq datasets in as few as 100 cells. We have shown that addition of carriers in FARP-ChIP-seq allows capture of low abundance chromatin without excessive PCR amplification, thereby greatly improves the fidelity and reproducibility of genome-wide chromatin mapping in small number of cells. However, a ~5-fold increase of sequencing depth is required to obtain sufficient reads of interest by FARP-ChIP-seq (Zheng *et al.*, 2015), which increases sequencing costs. The improved chromatin fragmentation by Atlantis DNase in aFARP-ChIP-seq greatly increased the recovery of low abundance chromatin compared to FARP-ChIP-seq. Indeed, we show that aFARP-ChIP-seq offers a ~3-fold increase of DNA reads compared to FARP-ChIP-seq at the same read depth for H3K4me3 in 500 mESCs. This increase in sequence recovery significantly reduces DNA reads needed and thus sequencing cost, thereby facilitating the use of carrier approach for ChIP-seq in different applications.

Not much effort has been devoted to multiplexing of ChIP-seq in small number of cells because of the difficulty in obtaining high quality and fidelity reads and because of the inconsistency of chromatin fragmentation by sonication, tagmentation, or MNase in different samples. We show that optimal chromatin fragmentation and high reproducibility is achieved by Atlantis dsDNase in a range of conditions and different cell number and types (Figue 1 and Figure 3-4). Therefore, aFARP-ChIP-seq offers a good opportunity to multiplexing ChIP-seq library preparations. This, coupled with the high-quality ChIP-seq data and reduced sequence need, should allow parallel epigenome mapping of different cell types isolated from different tissues without further amplification of cells or excessive rounds of PCR amplification of DNA fragments. The simplicity of aFARP-ChIP-seq workflow allows it to be easily established in any lab without specialized equipment, such as microfluidic devices used in MOWChIP-seq and single-cell ChIP-seq (Cao *et al.*, 2015; Rotem *et al.*, 2015). Considering the aFARP-ChIP-seq is fundamentally different from other high-sensitivity ChIP technologies, which rely on either excessive DNA amplification or chromatin indexing, we believe that it offers a viable alternative approach to further reduce read depth and cell number needed in a high throughout format.

By sorting the group 1 ILC subsets in mesenteric lymph node (mLN), spleen, small intestine, and liver from one mouse, we obtained high quality mapping of H3K4me3 and H3K27Ac in the NK and ILC1 cells. This should allow more accurate understanding of how the same kind of immune cell subsets differ in different organs in the same mouse. By obtaining genome wide maps of epigenome and chromatin binding sites for proteins in different cell types in different mouse, aFARP-ChIP-seq should also allow efficient comparisons of the same cell subsets in one organ to reveal how different environments and genotypes influence the genome features. For example, studies suggested that microenvironments in different tissues play important roles in shaping both gene expression and enhancer activities, resulting in tissue-specific identities of macrophages (Lavin *et al.*, 2014). Consistent with the notion that different tissue microenvironments influence tissue resident immune cells, our H3K4me3 MDS analyses reveal a clear separation of NK cells and ILC1s isolated from the spleen, small intestines, mLN, or liver in one mouse (Figure 3F).

We also observed pronounced epigenome differences in siIEL ILC1 isolated from small intestine’s intraepithelial compartment compared to the ILC1s from the other three tissues we analyzed, which is consistent with the recently reported transcriptome differences in the group I ILCs (Robinette *et al.*, 2015). Interestingly, we show that siIEL ILC1 exhibits H3K4me3 peaks on Eomes, a key lineage-determining transcription factor for NK cells. One explanation for this finding is that the siIEL ILC1 identified by cell-surface markers (NKp46+NK1.1+) is a heterogeneous population and it may be composed of both unidentified NK subsets and ILC1 lineage in the gut intraepithelial compartment. Alternatively, these siIEL ILC1 may exhibit lineage and functional plasticity due to the unique gut environment. Consistently, an ILC3-derived ILC1 population has been reported in the mouse gut under the influence of inflammatory stimuli (Li *et al.*, 2016). Importantly, NK cells could be converted into ILC1 by the tumor microenvironment-derived TGF-β signaling (Cortez *et al.*, 2017; Gao *et al.*, 2017), indicating that a fraction of siIEL ILC1 could also be derived from NK cells underlying the unique gut compartment. Identification of signaling molecules involved in the conversion of NK to siIEL ILC1 within the gut epithelial environment could be a key to understanding the functional plasticity of this unique ILC1 population. Given that gut microbiome is thought to exhibit remarkable impact on the regulatory landscape of ILCs (Gury-BenAri *et al.*, 2016a), aFARP-ChIP-seq should greatly aid further study of how ILCs differentially integrate signals from the microbial microenvironment in individual mice to generate phenotypic and functional differences, thereby influencing the health status of individuals.

The ability to achieve accurate map of active enhancer landscapes as revealed by H3K27Ac profiling in individual mice in this study should allow the identification of candidate genes that may function in specific cell types under different tissue microenvironments or external environments different mice experience. For example, compared to ILC1 from other peripheral tissues studied, the siIEL ILC1 displays a significant up-regulated H3K27Ac peaks for the gene Ahrr, which has been reported to play an important role for balancing colon inflammation (Brandstätter *et al.*, 2016). This suggests that Ahrr in siIEL ILC1 could play a similar role in the small intestines. Additionally, the H3K27Ac profiling allowed the identification of transcription factor binding signature enriched in siIEL ILC1. Deep mining of the H3K27Ac datasets in NK1 and ILC1 we mapped should allow the identification of additional candidate transcription factor binding in different tissues and cell subsets, which would facilitate further study of the regulatory logic underlying the developmental and homeostatic processes of group I ILCs.

Our proof-of-concept PU.1 profiling by aFARP-ChIP-seq using 1×10^4^ isolated splenic B cells demonstrate the broad usage of this method for genome-wide mapping of transcription factor binding. The requirement of only 1×10^4^ cells should allow successful mapping for majority of cell types of hematopoietic lineage in individual mice. This is especially important for the mapping of early hematopoietic progenitors because of limited cell number per mice. Additional optimization, including further optimizing digestion conditions and the use of improved ChIP grade antibodies, should allow the reduction of the number of cells needed, thereby further increasing the feasibility and success rate of discovering novel transcriptional regulatory network governing each stage of developmental process.

## Materials and methods

### Cell lines and animals

E14 mouse embryotic stem cells (mESCs) were cultured in DMEM with 15% fetal calf serum, penicillin/streptomycin, β-mercaptoethanol, L-glutamine, nonessential amino acids, recombinant leukemia inhibitory factor (1000 U/ml, Millipore).

C57BL/6L mice were obtained and maintained at the mouse facility of Carnegie Institution’s Embryology Department. Adult female mice (8 week of age) were used in all experiments. Mice were housed under a strict 12 hr light-dark cycle with food and water ad libitum. All mouse procedures in this study were in accordance with protocols approved by the Institutional Animal Care and Use Committee of the Carnegie Institution for Science.

### Antibodies

The following antibodies were obtained from Biolegend: Fluorescein isothiocyanate (FITC)-conjugated anti-NKp46 (clone 29A1.4), PE/Cy7-conjugated anti-NK1.1 (clone PK136), APC-conjugated anti-CD127 (IL-7Rα) (clone A7R34), PE/Cy7-conjugated anti-NK1.1 (clone PK136), PerCP/Cy5.5-conjugated anti-CD49b (clone HMa2), PerCP/Cy5.5-conjugated anti-mouse Ly-6A/E (Sca-1) (clone D7), PE anti-CD253(TRAIL) (clone N2B2), PE/Cy7-conjugated anti-mouse CD117 (clone c-Kit) (clone 2B8), FITC-conjugated anti-mouse CD25 (clone 3C7), Lineage mixed cocktail antibodies (Ter119, Gr1, CD11b, B220, CD3, CD4, CD8), PE-conjugated anti-F4/80 (clone BM8), APC-conjugated anti-CD11c (clone N418), and isotype-matched control monoclonal antibodies.

Antibody against histone H3 lysine 4 trimethylation (H3K4me3) (clone C42D8, #9751S) was from Cell Signaling. Antibodies against acetylated histone H3 lysine 27 (H3K27ac, #ab4729) and control mouse IgG were from Abcam. Antibodies against PU.1 (clone T-21, #sc-352X) were from Santa Cruz Biotechnology. The working concentrations of the above antibodies were used as recommended by the companies, unless otherwise specified below, or in the text and figure legends, in an assay-dependent manner.

### Cell identification, isolation, and flow cytometry

All cells from mouse tissues or organs were collected, stained and sorted according to the published standard protocol (Halim and Takei, 2014). Spleens, mesenteric lymph nodes, liver, and small intestine were extracted from C57BL/6J female mice and were dissociated into single cell suspensions by passing through 100µm Falcon cell strainer. After washing with FACS buffer (PBS with 0.3% BSA and 2 mM EDTA), cell suspension was incubated in red blood cell lysis solution (#R7757, Sigma) for 5 min on ice. For spleen and mesenteric lymph nodes, anti-CD3 (#130-095-130) and anti-CD19 (#130-052-201) microbeads (Miltenyi Biotec) were used to remove T cells and B cells, respectively, following the microbead guidelines. For liver, lymphocytes were enriched at the interface between a gradient of 40% and 80% Percoll in Hank’s balanced salt solution. For small intestine, the intestine was cut first longitudinally and then laterally into pieces of approximately 1-2 cm length in Petri dish. The tissues were transferred into 50 ml tube with 20 ml PBS containing 1mM EDTA. The sample was incubated at 37°C for 20 min under continuous 120 rpm rotation. Lymphocytes in small intestine were enriched at the interface between a gradient of 40% and 80% Percoll in PBS. The cells were then stained with DAPI, NKp46, NK1.1, CD127, CD49b, TRAIL, CD25, CD117, Sca-1, Lineage mixed cocktail antibodies (Ter119, Gr1, CD11b, B220, CD3, CD4, CD8), CD11c, I-A/I-E, F4/80 antibodies for 20 min at 4°C. After washing, cells were sorted with FACSAriaTM III cell sorter (BD Bioscience). All data were with FlowJo 9.3.2 software (Tree Star). The cell populations collected were identified as:

Spleen- and mesenteric lymph node-ILC1: CD3-, CD19-, NKp46+, NK1.1+, Spleen- and mesenteric lymph node-NK cells: CD3-, CD19-, NKp46+, NK1.1+, CD127-.

Liver-ILC1 cells: NKp46+, NK1.1+, CD49b-, TRAIL+.

Liver-NK cells: NKp46+, NK1.1+, CD49b+, TRAIL-.

siIEL-ILC1 cells: CD3-, CD19-, NKp46+, NK1.1+.

Spleen-B cells: CD3-, B220+, CD19+.

Spleen-T cells: CD3+, B220-. TCRβ+

### FARP-ChIP-seq

Standard FARP-ChIP-seq was performed according to the conditions previously from our lab (Zheng *et al.*, 2015). Briefly, 500 mESCs were mixed with ~5×10^8^ DH5α *E.coli* and fixed with 1% formaldehyde followed by quenching using 0.125M glycine. After washes, the sample mixtures were sonicated to obtain fragments with a 1/16-inch probe for 15 min at 3 watts by a tip sonicator (Misonix sonicator 3000). Protein G beads (#10004D, Life Technologies) and M-280 streptavidin beads (#11206D, Life Technologies) were pre-blocked with ~5×10^8^ fixed and sonicated *E.coli* lysate overnight at 4°C. After blocking, 5 ng carrier biotin-DNA was then coupled to 10 µl M-280 streptavidin beads. These treated protein G and streptavidin beads were combined and used to ChIP H3K4me3 in the sonicated *E.coli*+mESC lysates overnight at 4°C. After de-crosslinking, the precipitated genomic DNA and biotin-DNA were purified by AMPure XP beads (#A63881, Beckman Coulter). Library building steps were performed following Illumina True-Seq protocol with 0.25µM blocker oligo included at the final library amplification step.

### MNase-FARP-ChIP-seq

500 mESCs were mixed with ~1×10^8^ DH5α *E.coli* and fixed with 1% formaldehyde followed by quenching using 0.125M glycine. After fixation and washing, the mixtures were re-suspended directly in nuclear isolation buffer (#NUC101, Sigma). Chromatin was fragmented using 2U/μl MNase (#M2047 NEB) at 25°C for 5 minutes according to the published protocol (Brind’Amour *et al.*, 2015). ChIP, genomic DNA recovery, and sequencing library generation were performed following the FARP-ChIP-seq procedure.

### Atlantis-FARP-ChIP-seq (aFARP-ChIP-seq)

mESCs or sorted immune cells were mixed with ~1×10^8^ *E.coli* (DH5α) and crosslink by 1% formaldehyde and incubated for 8 min at room temperature with moderate shaking. After fixation, glycine was adding to the final concentration of 0.125 M and incubated for 5 min at room temperature to stop the crosslinking by quenching the free formaldehyde. After washing, the mixture was resuspended in the nuclear isolation buffer (#NUC101, Sigma). Then, the samples were digested with 0.5U/100ul Atalantis dsDNase (#E2030, Zymo Research) for 30 min at 37°C (20 min at 37°C for 100 mESCs). The reactions were stopped by 0.5 M EDTA. ChIP, genomic DNA recovery and sequencing library generation were performed following the standard FARP-ChIP-seq.

### ChIP sequencing and peak finding

ChIP sequencing was done on Illumina Nextseq-500 and pooled libraries were sequenced at a sequencing depth of ~15-20 million aligned reads per sample. Libraries were prepared in triplicates or duplicates. Reads were mapped to the mouse genome mm9 using the ‘bowtie’ program with -v 2 parameter. Only tags that uniquely mapped to the genome were used for further analysis. ChIP-seq peaks were called using the MACS program (Zhang *et al.*, 2008) with default parameters.

### Promoter correlation, whole-genome correlation, and ROC analysis

log2 of H3K4me3 enrichment were plotted for all TSS (2kb up and downstream of TSS) to generate the correlation plots. For the H3K4me3 ROC analysis, we used top 25,000 2-kb windows as “True” to mimic the ~25,000 peaks in benchmark dataset. Then by adjusting the “threshold” to include more top 2-kb windows in the test data, we can calculate the true-positive rate and false-positive rate to get the ROC curve.

### Analyses of ChIP-seq of transcription factor binding in splenic B cells

PU.1 peaks of B cells from mouse spleen are called by MACS with p-value threshold of 10^−5^. The motifs was identified by using Homer (Heinz *et al.*, 2010) to call motifs on PU.1 peak regions.

**Supplementary Figure legends**

**Figure S1.**
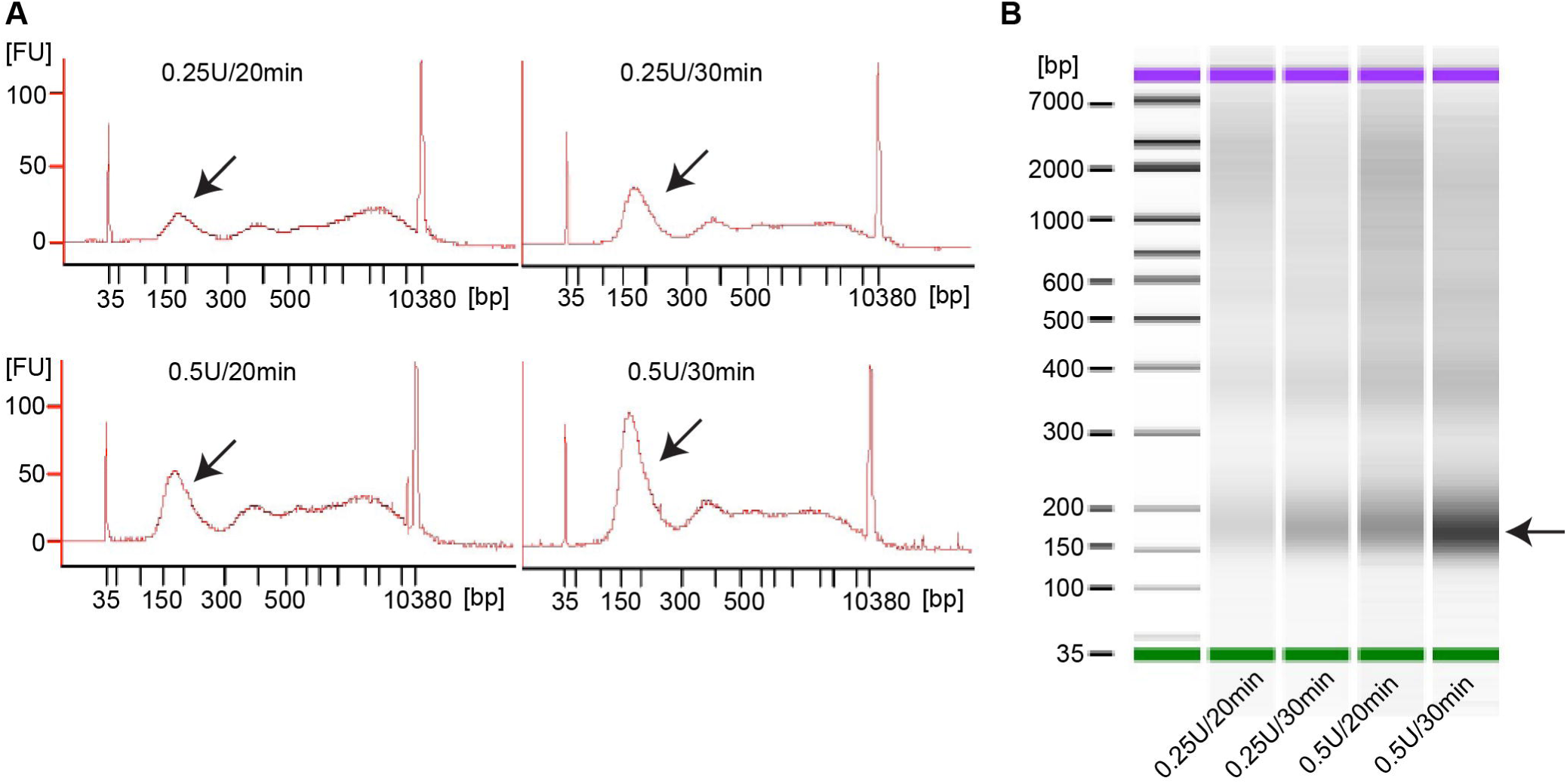
Analyses of chromatin fragmentation conditions by Atlantis dsDNase. (A-B) Bioanalyzer plots (A) and gel analyses (B) showing the efficiency of chromatin fragmentation using 1×10^5^ mESCs by Atlantis dsDNAse with the indicated enzyme unit and digestion time. Arrows mark the position of mono-nucleosome.

**Figure S2.**
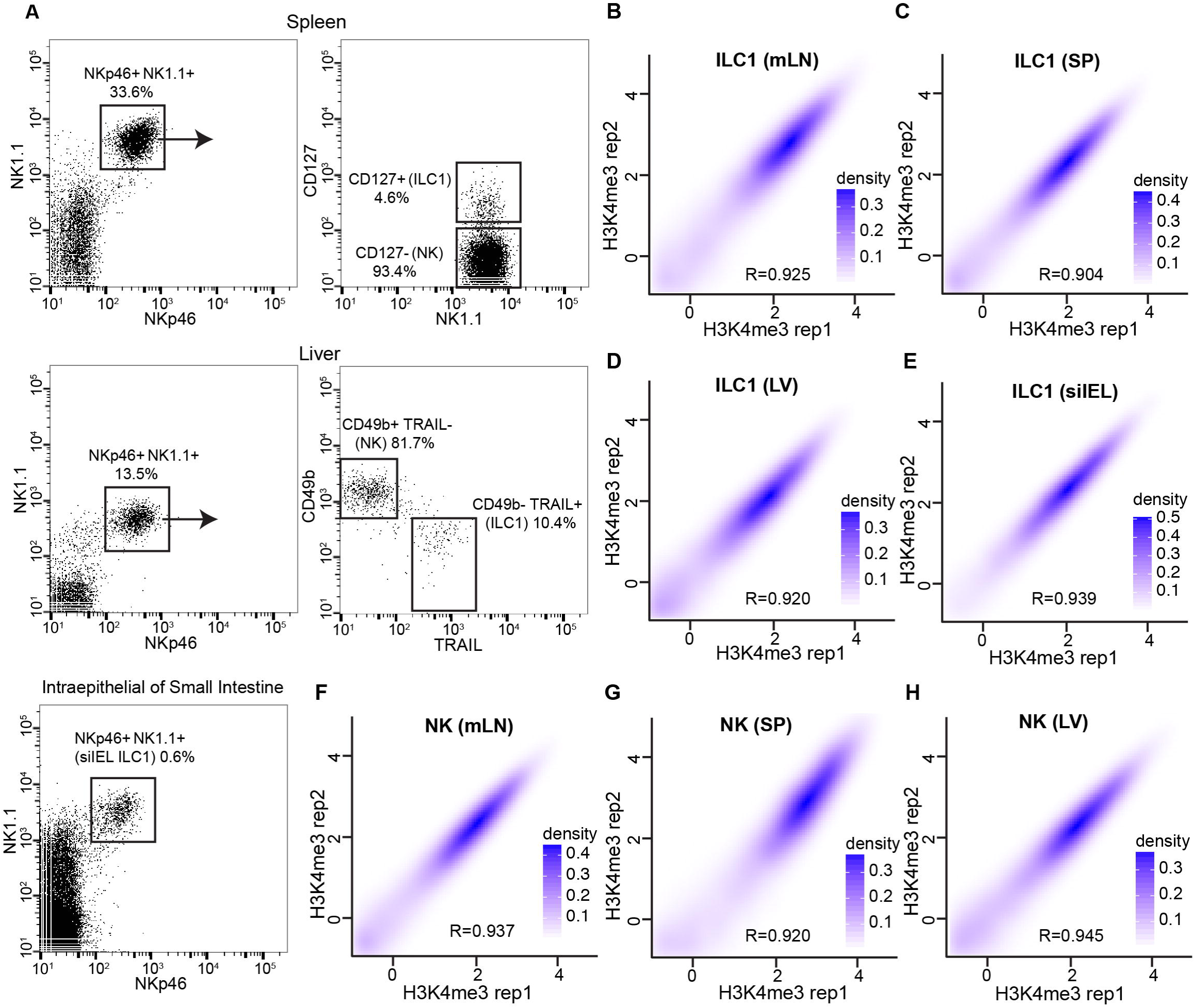
aFARP-ChIP-seq of H3K4me3 in group 1 ILCs from different tissues. (A) FACS plots with the markers and gating used for sorting NK and ILC1 cells from spleen (SP) and liver (LV) or ILC1 from intraepithelial of small intestine (siIEL). (B-E) Contour plots between two biological replicates of aFARP-ChIP-seq of H3K4me3 in ILC1 in mLN (B), SP (C), LV (D), and siIEL (E). (F-H) Contour plots between two biological replicates of aFARP-ChIP-seq of H3K4me3 in NK cells isolated from mLN (F), SP (G), and LV (H).

**Figure S3.**
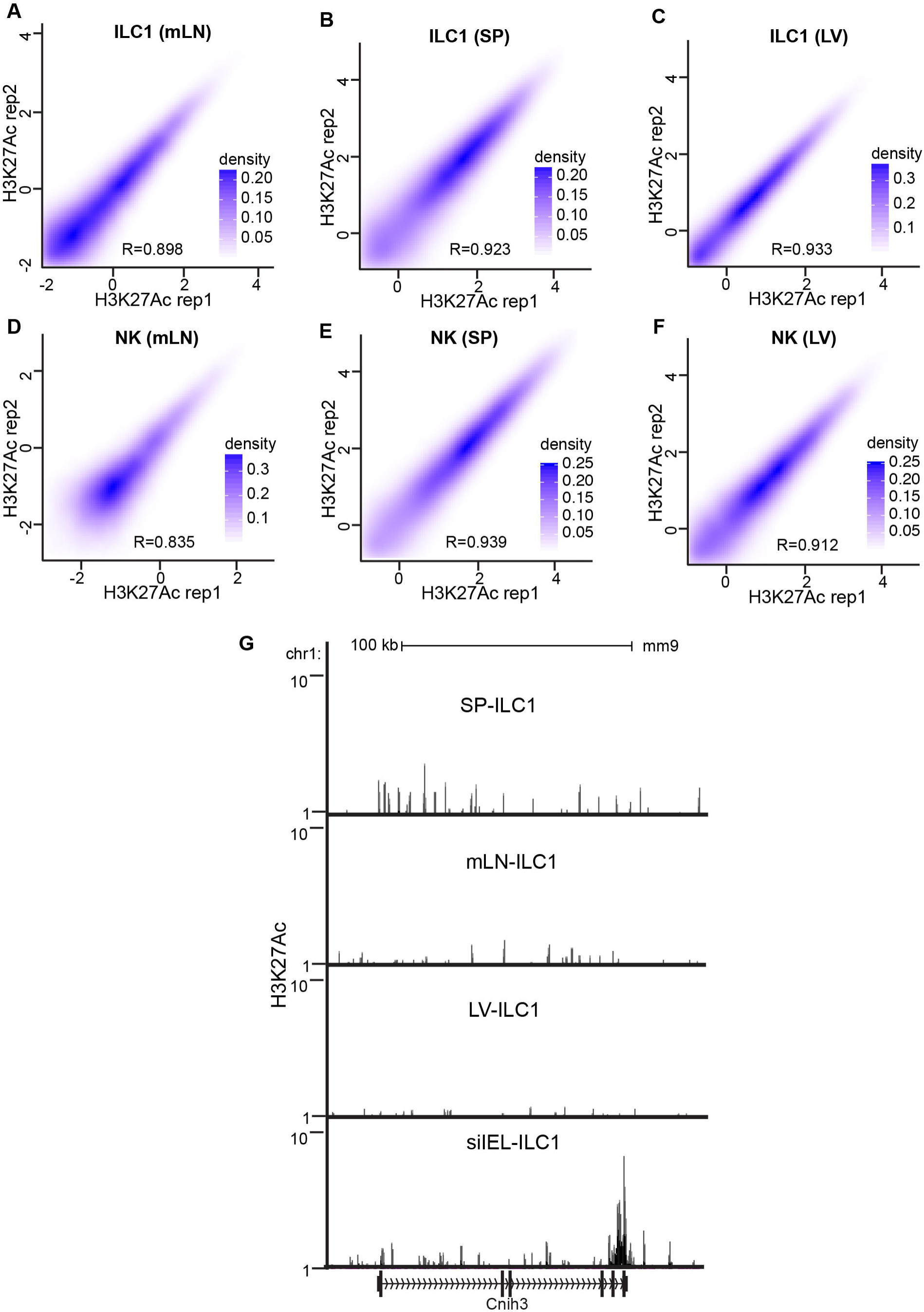
aFARP-ChIP-seq of H3K27Ac in group 1 ILCs from different tissues. (A-C) Contour plots between two biological replicates of aFARP-ChIP-seq of H3K27Ac in ILC1 in mLN (A), SP (B), and LV (C). (D-F) Contour plots between two biological replicates of aFARP-ChIP-seq of H3K27Ac in NK cells in mLN (D), SP (E) and LV (F). (H) Browser view of H3K27Ac map at Cnih3 in ILC1s from the indicated tissues.

**Figure S4.**
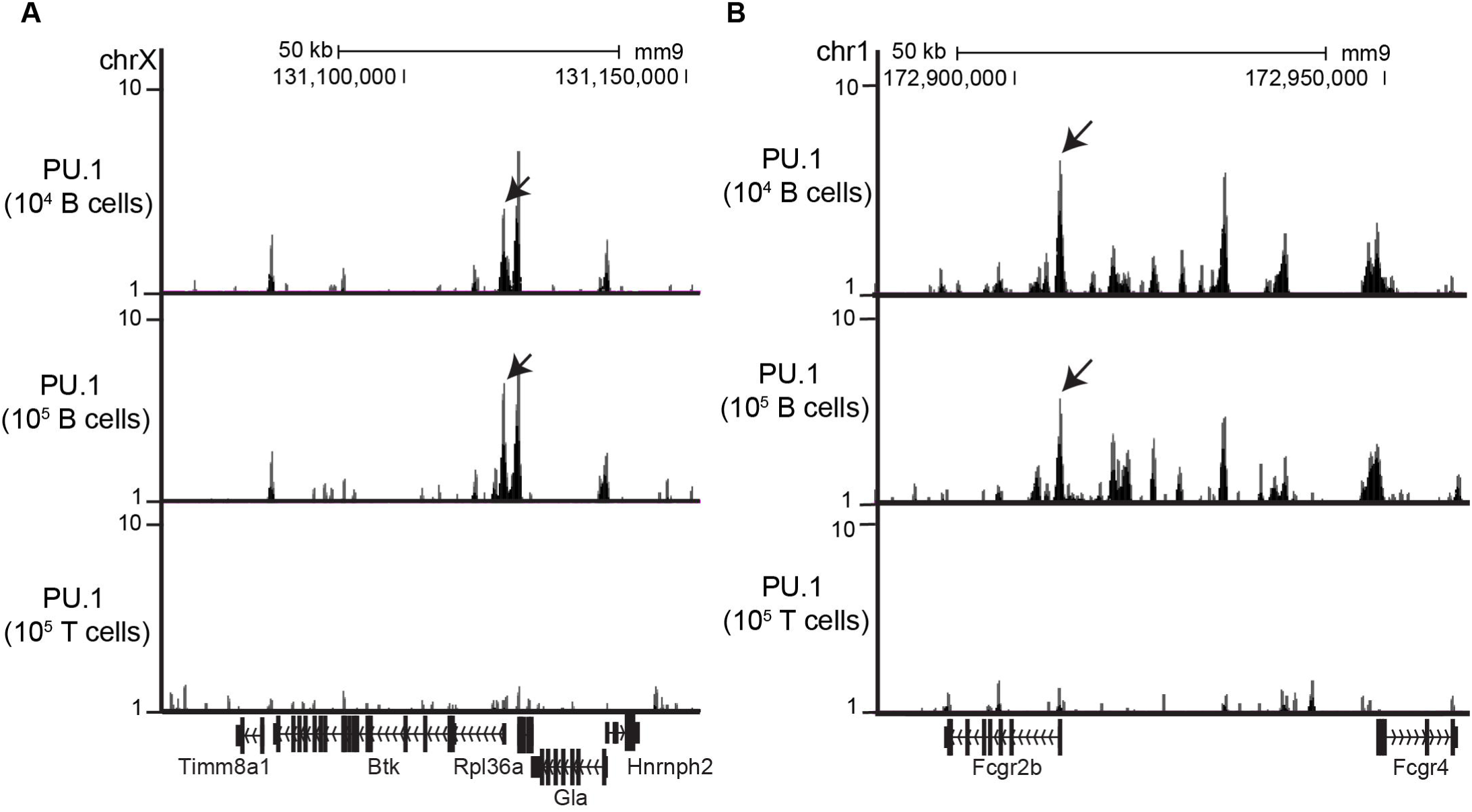
aFARP-ChIP-seq of PU.1 profiling in splenic B cells. (A-B) Normalized profiles of PU.1 in a 75 kb region surrounding the Btk (A) or Fcgr2b (B) loci obtained with aFARP-ChIP-seq in isolated splenic B and T cells. The input cell numbers are indicated on the y axis. Arrows mark peaks in the Btk (A) or Fcgr2b (B) locus.

**Supplementary Table legends**

**Table S1**. Peaks called from H3K4me3 mapping of ILC1 and NK cells from indicated tissues. Related to Figure 3. Replicate datasets are pooled before peak calling

**Table S2**. Peaks called from H3K27ac mapping of ILC1 and NK cells from indicated tissues. Related to Figure 4. Replicate datasets are pooled before peak calling.

**Table S3**. PU.1 peaks called from mapping of B cells from mouse spleen. Related to Figure 5.

## References

Adli, M., and Bernstein, B.E. (2011). Whole-genome chromatin profiling from limited numbers of cells using nano-ChIP-seq. Nat Protoc 6, 1656–1668.

Adli, M., Zhu, J., and Bernstein, B.E. (2010). Genome-wide chromatin maps derived from limited numbers of hematopoietic progenitors. Nat Methods 7, 615–618.

Brandstätter, O., Schanz, O., Vorac, J., König, J., Mori, T., Maruyama, T., Korkowski, M., Haarmann-Stemmann, T., von Smolinski, D., Schultze, J.L., Abel, J., Esser, C., Takeyama, H., Weighardt, H., and Förster, I. (2016). Balancing intestinal and systemic inflammation through cell type-specific expression of the aryl hydrocarbon receptor repressor. Sci Rep 6, 26091.

Brind’Amour, J., Liu, S., Hudson, M., Chen, C., Karimi, M.M., and Lorincz, M.C. (2015). An ultra-low-input native ChIP-seq protocol for genome-wide profiling of rare cell populations. Nat Commun 6, 6033.

Cao, Z., Chen, C., He, B., Tan, K., and Lu, C. (2015). A microfluidic device for epigenomic profiling using 100 cells. Nat Methods 12, 959–962.

Cortez, V.S., Cervantes-Barragan, L., Robinette, M.L., Bando, J.K., Wang, Y., Geiger, T.L., Gilfillan, S., Fuchs, A., Vivier, E., Sun, J.C., Cella, M., and Colonna, M. (2016). Transforming Growth Factor-β Signaling Guides the Differentiation of Innate Lymphoid Cells in Salivary Glands. Immunity 44, 1127–1139.

Cortez, V.S., and Colonna, M. (2016). Diversity and function of group 1 innate lymphoid cells. Immunol Lett 179, 19–24.

Cortez, V.S., Ulland, T.K., Cervantes-Barragan, L., Bando, J.K., Robinette, M.L., Wang, Q., White, A.J., Gilfillan, S., Cella, M., and Colonna, M. (2017). SMAD4 impedes the conversion of NK cells into ILC1-like cells by curtailing non-canonical TGF-β signaling. Nat Immunol 18, 995–1003.

Creyghton, M.P., Cheng, A.W., Welstead, G.G., Kooistra, T., Carey, B.W., Steine, E.J., Hanna, J., Lodato, M.A., Frampton, G.M., Sharp, P.A., Boyer, L.A., Young, R.A., and Jaenisch, R. (2010). Histone H3K27ac separates active from poised enhancers and predicts developmental state. Proc Natl Acad Sci U S A 107, 21931–21936.

Daussy, C., Faure, F., Mayol, K., Viel, S., Gasteiger, G., Charrier, E., Bienvenu, J., Henry, T., Debien, E., Hasan, U.A., Marvel, J., Yoh, K., Takahashi, S., Prinz, I., de Bernard, S., Buffat, L., and Walzer, T. (2014). T-bet and Eomes instruct the development of two distinct natural killer cell lineages in the liver and in the bone marrow. J Exp Med 211, 563–577.

Diefenbach, A., Colonna, M., and Koyasu, S. (2014). Development, differentiation, and diversity of innate lymphoid cells. Immunity 41, 354–365.

Eberl, G., Colonna, M., Di Santo, J.P., and McKenzie, A.N. (2015). Innate lymphoid cells. Innate lymphoid cells: a new paradigm in immunology. Science 348, aaa6566.

Fuchs, A., Vermi, W., Lee, J.S., Lonardi, S., Gilfillan, S., Newberry, R.D., Cella, M., and Colonna, M. (2013). Intraepithelial type 1 innate lymphoid cells are a unique subset of IL-12-and IL-15-responsive IFN-γ-producing cells. Immunity 38, 769–781.

Furey, T.S. (2012). ChIP-seq and beyond: new and improved methodologies to detect and characterize protein-DNA interactions. Nat Rev Genet 13, 840–852.

Gao, Y., Souza-Fonseca-Guimaraes, F., Bald, T., Ng, S.S., Young, A., Ngiow, S.F., Rautela, J., Straube, J., Waddell, N., Blake, S.J., Yan, J., Bartholin, L., Lee, J.S., Vivier, E., Takeda, K., Messaoudene, M., Zitvogel, L., Teng, M.W.L., Belz, G.T., Engwerda, C.R., Huntington, N.D., Nakamura, K., Hölzel, M., and Smyth, M.J. (2017). Tumor immunoevasion by the conversion of effector NK cells into type 1 innate lymphoid cells. Nat Immunol 18, 1004–1015.

Gordon, S.M., Chaix, J., Rupp, L.J., Wu, J., Madera, S., Sun, J.C., Lindsten, T., and Reiner, S.L. (2012). The transcription factors T-bet and Eomes control key checkpoints of natural killer cell maturation. Immunity 36, 55–67.

Gury-BenAri, M., Thaiss, C.A., Serafini, N., Winter, D.R., Giladi, A., Lara-Astiaso, D., Levy, M., Salame, T.M., Weiner, A., David, E., Shapiro, H., Dori-Bachash, M., Pevsner-Fischer, M., Lorenzo-Vivas, E., Keren-Shaul, H., Paul, F., Harmelin, A., Eberl, G., Itzkovitz, S., Tanay, A., Di Santo, J.P., Elinav, E., and Amit, I. (2016a). The Spectrum and Regulatory Landscape of Intestinal Innate Lymphoid Cells Are Shaped by the Microbiome. Cell 166, 1231–1246 e1213.

Gury-BenAri, M., Thaiss, C.A., Serafini, N., Winter, D.R., Giladi, A., Lara-Astiaso, D., Levy, M., Salame, T.M., Weiner, A., David, E., Shapiro, H., Dori-Bachash, M., Pevsner-Fischer, M., Lorenzo-Vivas, E., Keren-Shaul, H., Paul, F., Harmelin, A., Eberl, G., Itzkovitz, S., Tanay, A., Di Santo, J.P., Elinav, E., and Amit, I. (2016b). The Spectrum and Regulatory Landscape of Intestinal Innate Lymphoid Cells Are Shaped by the Microbiome. Cell 166, 1231–1246.e1213.

Halim, T.Y., and Takei, F. (2014). Isolation and characterization of mouse innate lymphoid cells. Curr Protoc Immunol 106, 3.25.21–13.

Hallikas, O., Palin, K., Sinjushina, N., Rautiainen, R., Partanen, J., Ukkonen, E., and Taipale, J. (2006). Genome-wide prediction of mammalian enhancers based on analysis of transcription-factor binding affinity. Cell 124, 47–59.

Heinz, S., Benner, C., Spann, N., Bertolino, E., Lin, Y.C., Laslo, P., Cheng, J.X., Murre, C., Singh, H., and Glass, C.K. (2010). Simple combinations of lineage-determining transcription factors prime cis-regulatory elements required for macrophage and B cell identities. Mol Cell 38, 576–589.

Heinz, S., Romanoski, C.E., Benner, C., and Glass, C.K. (2015). The selection and function of cell type-specific enhancers. Nat Rev Mol Cell Biol b16, 144–154.

Jia, J., Zheng, X., Hu, G., Cui, K., Zhang, J., Zhang, A., Jiang, H., Lu, B., Yates, J., Liu, C., Zhao, K., and Zheng, Y. (2012). Regulation of pluripotency and self-renewal of ESCs through epigenetic-threshold modulation and mRNA pruning. Cell 151, 576–589.

Jiao, Y., Huntington, N.D., Belz, G.T., and Seillet, C. (2016). Type 1 Innate Lymphoid Cell Biology: Lessons Learnt from Natural Killer Cells. Front Immunol 7, 426.

Klemsz, M.J., McKercher, S.R., Celada, A., Van Beveren, C., and Maki, R.A. (1990). The macrophage and B cell-specific transcription factor PU.1 is related to the ets oncogene. Cell 61, 113–124.

Lara-Astiaso, D., Weiner, A., Lorenzo-Vivas, E., Zaretsky, I., Jaitin, D.A., David, E., Keren-Shaul, H., Mildner, A., Winter, D., Jung, S., Friedman, N., and Amit, I. (2014). Immunogenetics. Chromatin state dynamics during blood formation. Science 345, 943–949.

Lavin, Y., Winter, D., Blecher-Gonen, R., David, E., Keren-Shaul, H., Merad, M., Jung, S., and Amit, I. (2014). Tissue-resident macrophage enhancer landscapes are shaped by the local microenvironment. Cell 159, 1312–1326.

Li, J., Doty, A., and Glover, S.C. (2016). Aryl hydrocarbon receptor signaling involves in the human intestinal ILC3/ILC1 conversion in the inflamed terminal ileum of Crohn’s disease patients. Inflamm Cell Signal 3.

McKenzie, A.N.J., Spits, H., and Eberl, G. (2014). Innate lymphoid cells in inflammation and immunity. Immunity 41, 366–374.

Ng, J.H., Kumar, V., Muratani, M., Kraus, P., Yeo, J.C., Yaw, L.P., Xue, K., Lufkin, T., Prabhakar, S., and Ng, H.H. (2013). In vivo epigenomic profiling of germ cells reveals germ cell molecular signatures. Dev Cell 24, 324–333.

Ouyang, Z., Zhou, Q., and Wong, W.H. (2009). ChIP-Seq of transcription factors predicts absolute and differential gene expression in embryonic stem cells. Proc Natl Acad Sci U S A 106, 21521–21526.

Park, P.J. (2009). ChIP-seq: advantages and challenges of a maturing technology. Nat Rev Genet 10, 669–680.

Rankin, L., Groom, J., Mielke, L.A., Seillet, C., and Belz, G.T. (2013). Diversity, function, and transcriptional regulation of gut innate lymphocytes. Front Immunol 4, 22.

Robinette, M.L., Fuchs, A., Cortez, V.S., Lee, J.S., Wang, Y., Durum, S.K., Gilfillan, S., Colonna, M., and Consortium, I.G. (2015). Transcriptional programs define molecular characteristics of innate lymphoid cell classes and subsets. Nat Immunol 16, 306–317.

Rotem, A., Ram, O., Shoresh, N., Sperling, R.A., Goren, A., Weitz, D.A., and Bernstein, B.E. (2015). Single-cell ChIP-seq reveals cell subpopulations defined by chromatin state. Nat Biotechnol 33, 1165–1172.

Schmidl, C., Rendeiro, A.F., Sheffield, N.C., and Bock, C. (2015). ChIPmentation: fast, robust, low-input ChIP-seq for histones and transcription factors. Nat Methods 12, 963–965.

Schweitzer, B.L., Huang, K.J., Kamath, M.B., Emelyanov, A.V., Birshtein, B.K., and DeKoter, R.P. (2006). Spi-C has opposing effects to PU.1 on gene expression in progenitor B cells. J Immunol 177, 2195–2207.

Sciumè, G., Shih, H.Y., Mikami, Y., and O’Shea, J.J. (2017). Epigenomic Views of Innate Lymphoid Cells. Front Immunol 8, 1579.

Shankaranarayanan, P., Mendoza-Parra, M.A., Walia, M., Wang, L., Li, N., Trindade, L.M., and Gronemeyer, H. (2011). Single-tube linear DNA amplification (LinDA) for robust ChIP-seq. Nat Methods 8, 565–567.

Shih, H.Y., Sciumè, G., Mikami, Y., Guo, L., Sun, H.W., Brooks, S.R., Urban, J.F., Davis, F.P., Kanno, Y., and O’Shea, J.J. (2016). Developmental Acquisition of Regulomes Underlies Innate Lymphoid Cell Functionality. Cell 165, 1120–1133.

Solomon, L.A., Li, S.K., Piskorz, J., Xu, L.S., and DeKoter, R.P. (2015). Genome-wide comparison of PU.1 and Spi-B binding sites in a mouse B lymphoma cell line. BMC Genomics 16, 76.

Spitz, F., and Furlong, E.E. (2012). Transcription factors: from enhancer binding to developmental control. Nat Rev Genet 13, 613–626.

Stathopulos, P.B., Scholz, G.A., Hwang, Y.M., Rumfeldt, J.A., Lepock, J.R., and Meiering, E.M. (2004). Sonication of proteins causes formation of aggregates that resemble amyloid. Protein Sci 13, 3017–3027.

Tanriver, Y., and Diefenbach, A. (2014). Transcription factors controlling development and function of innate lymphoid cells. Int Immunol 26, 119–128.

Valouev, A., Johnson, D.S., Sundquist, A., Medina, C., Anton, E., Batzoglou, S., Myers, R.M., and Sidow, A. (2008). Genome-wide analysis of transcription factor binding sites based on ChIP-Seq data. Nat Methods 5, 829–834.

van Galen, P., Viny, A.D., Ram, O., Ryan, R.J., Cotton, M.J., Donohue, L., Sievers, C., Drier, Y., Liau, B.B., Gillespie, S.M., Carroll, K.M., Cross, M.B., Levine, R.L., and Bernstein, B.E. (2016). A Multiplexed System for Quantitative Comparisons of Chromatin Landscapes. Mol Cell 61, 170–180.

Xu, L.S., Sokalski, K.M., Hotke, K., Christie, D.A., Zarnett, O., Piskorz, J., Thillainadesan, G., Torchia, J., and DeKoter, R.P. (2012). Regulation of B cell linker protein transcription by PU.1 and Spi-B in murine B cell acute lymphoblastic leukemia. J Immunol 189, 3347–3354.

Zhang, J., Marotel, M., Fauteux-Daniel, S., Mathieu, A.L., Viel, S., Marçais, A., and Walzer, T. (2018). T-bet and Eomes govern differentiation and function of mouse and human NK cells and ILC1. Eur J Immunol 48, 738–750.

Zhang, Y., Liu, T., Meyer, C.A., Eeckhoute, J., Johnson, D.S., Bernstein, B.E., Nusbaum, C., Myers, R.M., Brown, M., Li, W., and Liu, X.S. (2008). Model-based analysis of ChIP-Seq (MACS). Genome Biol 9, R137.

Zheng, X., Yue, S., Chen, H., Weber, B., Jia, J., and Zheng, Y. (2015). Low-Cell-Number Epigenome Profiling Aids the Study of Lens Aging and Hematopoiesis. Cell Rep 13, 1505–1518.

